# lydemapr: an R package to track the spread of the invasive Spotted Lanternfly (*Lycorma delicatula*, White 1845) (Hemiptera, Fulgoridae) in the United States

**DOI:** 10.1101/2023.01.27.525992

**Authors:** Sebastiano De Bona, Lawrence Barringer, Paul Kurtz, Jay Losiewicz, Gregory R. Parra, Matthew R. Helmus

## Abstract

A crucial asset in the management of invasive species is the open-access sharing of data on the range of invaders and the progression of their spread. Such data should be current, comprehensive, consistent, and standardized, to support reproducible and comparable forecasting efforts among multiple researchers and managers. Here, we present the `lydemapr` R package containing spatiotemporal data and mapping functions to visualize the current spread of the spotted lanternfly (*Lycorma delicatula*, White 1841) in the Western Hemisphere. The spotted lanternfly is a forest and agricultural pest in the eastern Mid-Atlantic region of the U.S., where it was first discovered in 2014. As of 2022, it has been found in 12 states according to state and federal departments of agriculture. However, the lack of easily accessible, fine-scale data on its spread hampers research and management efforts. We obtained multiple memoranda-of-understanding from several agencies and citizen-science projects, gaining access to their internal data on spotted lanternfly point observations. We then cleaned, harmonized, anonymized, and combined the individual data sources into a single comprehensive dataset. The resulting dataset contains spatial data gridded at the 1 km^2^ resolution, with yearly information on the presence/absence of spotted lanternflies, establishment status, and population density across 658,392 observations. The `lydemapr` package will aid researchers, managers, and the public in their understanding, modeling, and managing of the spread of this invasive pest.

## Introduction

Due to the globalization of trade, and the homogenization of urban and suburban habitats, the accidental introduction and establishment of invasive species is ever more likely (Hulme, 2009). When establishment goes undetected and eradication becomes less viable, the goal should be to mitigate the negative effects generated by invasive species (Diagne et al., 2020; Fantle-Lepczyk et al., 2022; Leroy et al., 2022). In doing so, one of the main challenges is tracking the spread of established invasive alien species so that control measures to slow spread, reduce impact, and conserve biodiversity can be effectively enacted (Robertson et al., 2020). High quality data on past and present spread of invasives is key to model invasive spread accurately enough to provide robust forecasts on which to base management decisions.

A multitude of modelling techniques to forecast spread is available to researchers (Clark et al., 2003; Fisher, 1937; Higgins & Richardson, 1996; Hudgins et al., 2017; Jongejans et al., 2011; Kot et al., 1996; Neubert et al., 2000; Rodrigues & Johnstone, 2014; Skellam, 1951; Travis & Dytham, 2002). Despite different assumptions and approaches to the modeling itself, many of these techniques rely on longitudinal, spatially-explicit data on the occurrence or density of the spreading invasive species. Different models need to be built upon the same standardized data for comparisons between models to reflect genuine differences in model assumptions and to highlight what biological aspects of spread are crucial to manage. In addition, building models on the same data provides a more solid ground to combine them into ensemble models, which offer a higher degree of reliability than a single model (Araújo & New, 2007). However, there are three hurdles that must be overcome before such standardized data for modeling be made available.

The first hurdle that must be overcome when developing a standardized data set on invasive spread is to develop relationships with the agencies, institutions, and citizen science projects collecting data on the invasive of interest. For pests with negative impact on agricultural activity or forest habitats, local agencies, state departments, and research institutions associated with the species first discovery are likely to operate data collection. If the pest is spreading across geopolitical boundaries, multiple organizations with different jurisdictions and area of operation are likely to collect field data. In addition, easy-to-identify pests are likely to attract public attention and involvement, fostering the collection of citizen science data (Catlin-Groves, 2012; Dickinson et al., 2010; Johnson et al., 2020; Kobori et al., 2016; Norman-Burgdolf & Rieske, 2021, 2021; Santaoja, 2022; Sullivan et al., 2014). Obtaining access to the data often requires directly contacting the maintainer of the dataset in the relevant institution and getting memoranda-of-understanding to use the data once shared. Each agency will follow unique data sharing agreements, which need to be discussed in-depth at this stage.

Once the data is obtained, the heterogeneity of the data collection protocols adopted by different agencies requires several additional steps to harmonize the survey results before they can be combined into a single dataset. This second hurdle is often the most time consuming and requires a high degree of skill in data handling and management. Non-standardized data collection demands an in-depth understanding of the collection protocols used in order to match the information collected across different surveys. For this reason, harmonization often demands an active collaboration with the agencies that collected the data, to ensure the data is interpreted correctly, especially when surveys lack metadata.

The third hurdle is essential, yet not often acknowledged: data anonymization. Calls to make scientific knowledge more accessible and transparent have pushed ecological data to be published alongside many scientific papers (Reichman et al., 2011). This process is paramount to improve collaboration and repeatability of scientific studies, although some limitations need to occur to ensure sharing open access data is done safely (Lindenmayer & Scheele, 2017; Lunghi et al., 2019). One such limitation concerns data at high spatial resolution, the publication of which could infringe upon individual privacy and personal interests (Zipper et al., 2019). Because of this, invasive spread data needs to be carefully and fully anonymized to ensure stakeholders are protected and served. This is especially true when knowledge on the infested state of a property could cause its value to decrease, or the value of the goods produced to be affected (Kovacs et al., 2011; Zhang & Boyle, 2010). Anonymization practices include the removal of personal information, as well as data handling that reduces the spatial resolution to an optimal compromise between conveying relevant information and safeguarding privacy.

The spotted lanternfly (*Lycorma delicatula*, White 1845; often referred to as SLF in the literature) was first discovered in the United States in Berks County, Pennsylvania, in 2014 (Barringer et al., 2015; Dara et al., 2015), and has since spread to 12 states across the Mid-Atlantic and Midwestern United States (NYIPM, 2022; Urban et al., 2021). This phloem-feeding planthopper is native to China and was likely introduced accidentally via a shipment of landscaping materials (Urban, 2020). The spotted lanternfly is known to feed on over 100 species of plants (Barringer & Ciafré, 2020; Huron & Helmus, 2022; Murman et al., 2020) and poses a major economic burden on viticulture as it feeds on grapevine reducing total yield and plant vigor (Urban, 2020). State agencies and the United States Department of Agriculture (USDA) have collected large amounts of data on the spread of this pest through field surveys. In addition, given the species is easily recognized and hard to misidentify, a broad public campaign to educate the public has promoted the collection of citizen science data. Data is collected through individual use of well-established applications such as iNaturalist, which allow for users to record geo-referenced observations of wildlife sightings, as well as through the use of applications developed *ad hoc* by State departments of agriculture to collect data on the spotted lanternfly. Given the variety of sources, and the refinement of protocols for data collection, the data on this species is heavily heterogeneous. Currently, any research team analyzing the spread of the pest has to invest a significant amount of time processing the data before using it in model construction and validation (Cook et al., 2021; e.g. Huron et al., 2022; Jones et al., 2022; Ramirez et al., 2022; Wakie et al., 2020).

Here, we describe the R package `lydemapr` (**Ly**corma **de**licatula **map**ping in **R**), containing an up-to-date, fully anonymized, and regularly refined longitudinal, spatially-explicit dataset of spotted lanternfly records throughout the United States since its first discovery. The dataset includes information derived from field surveys and citizen-science observations, and reports observed presence/absence of this invasive species in surveyed areas, as well as the presence of established populations, and estimates of population density. In addition, the package contains tools to visualize the data by mapping it, and to obtain summary tables of the dataset. The goal of this package is to provide a baseline for future modeling efforts to forecast the spread of the spotted lanternfly, and to foster more effective collaboration between agencies and researchers. The `lydemapr` package was fully developed in R (R Core Team, 2021) and is available as an online repository at https://github.com/ieco-lab/lydemapr.

### Data and metadata

The dataset contained in the package represents an anonymized and condensed comprehensive record of data collected by several federal agencies, state agencies, and citizen-science projects on the presence, establishment, and population density of spotted lanternfly in the United States (Figure 1). Sources include the Departments of Agriculture for the states of Pennsylvania, Delaware, Indiana, Maryland, and New Jersey; the New York State Department of Agriculture and Markets; the Virginia Department of Agriculture and Consumer Services; the Virginia Polytechnic Institute and State University; the United States Department of Agriculture; and public reporting from iNaturalist. The field data was collected through a variety of methods, including surveys aiming to estimate establishment status and spotted lanternfly population density, control actions to manage population through egg mass destruction and trapping, and citizen science data collected through self-reporting or direct involvement through research projects. Self-reporting tools include two separate platforms developed by the Pennsylvania Department of Agriculture (PDA) in association with Penn State University (PSU) and the New Jersey Department of Agriculture (NJDA). In addition, we included data collected through an independent citizen-science projects of limited duration run by the Virginia Polytechnic Institute and State University and the Virginia Cooperative Extension.

**Figure 1.**
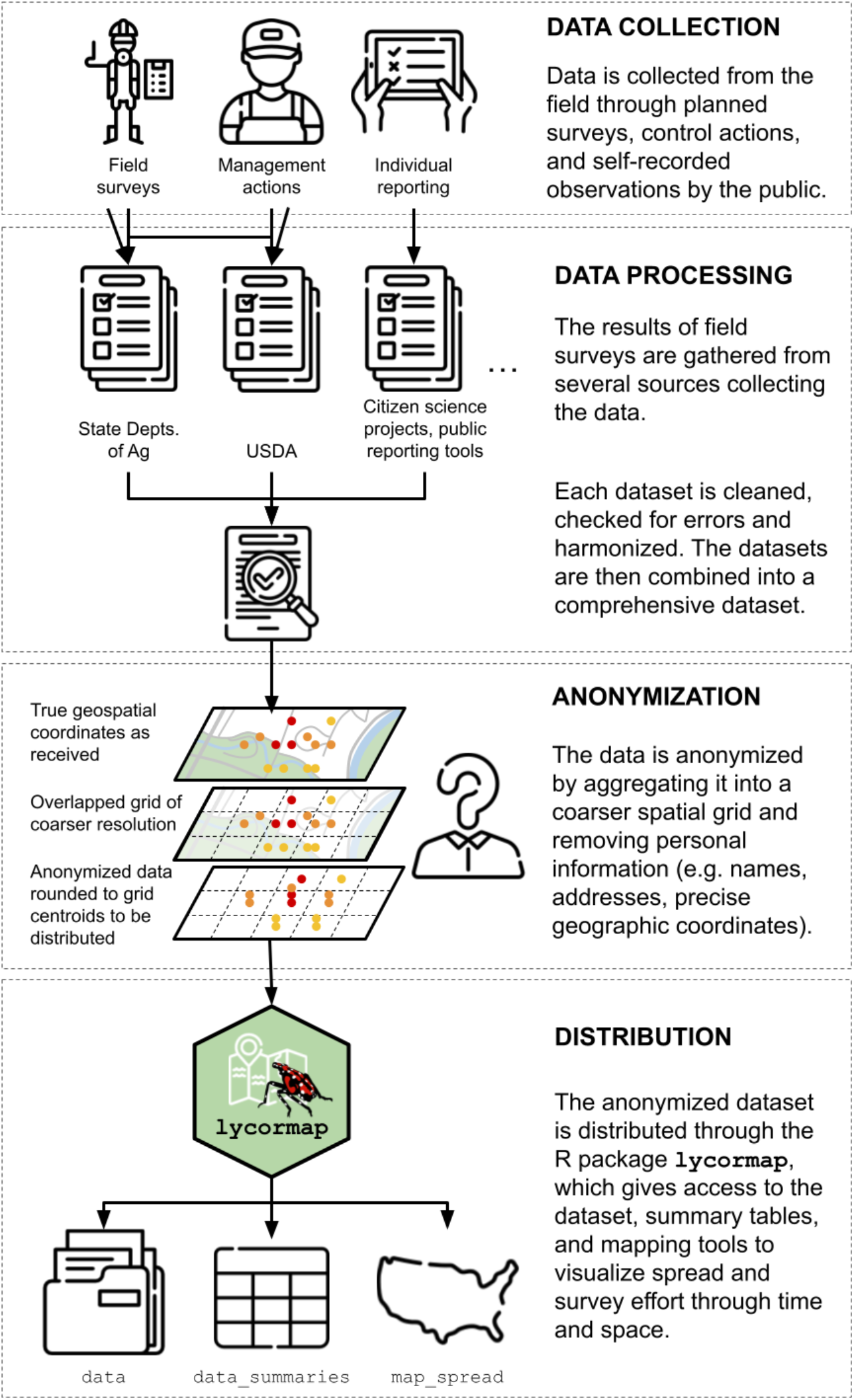
Conceptual graph describing the process leading to the distribution of the R package `lydemapr`. Data is collected by individual sources through multiple surveying processes. The dataset compiled this way are gathered from the sources and individually processed, then combined. The resulting comprehensive dataset is anonymized through both a censoring step and a spatial transformation. Any column containing information on individual identities and business names is removed. In addition, any information on the individual collecting the data is removed. For the spatial transformation, latitude and longitude of individual survey points are rounded to the centroids of a 1km^2^ resolution grid. The aggregated and anonymized dataset is distributed through the package, together with functions to visualize it.

At the date of publication, the aggregated and anonymized dataset contains 658,392 individual observations pertaining to 61,715 point-locations throughout the United States collected between 2014 and 2022. These 61,715 point-locations represent centroids of a 1 km^2^ grid at which the geospatial data was aggregated for anonymization. The exact latitude and longitude of each survey contained in the geospatial data collected by the sources are rounded to the coordinates of the centroids. This approach, while removing the ability to derive property-level information from the dataset, allows us to distribute survey-level information the data user can summarize as it best fits their needs. All variables containing traceable information regarding personal names, business names, contact information, and comments, were also removed from the dataset.

#### Variables included

- *source*: character variable defining in broad terms the source of the data. “inat” for data obtained from iNaturalist, “PA” from data originating from the Pennsylvania Dept. of Agriculture’s surveying and management effort, “prt” for data collected through public reporting platforms, “states” for data collected by state-level agencies other than PDA, “USDA” for data provided by the United States Dept of Agriculture. Note: the data originating from the Pennsylvania Dept. of Agriculture is kept separate from data collected by other states, as Pennsylvania was the state were the first introduction was detected. Because of this, initial surveying efforts were led by this state, which collected the largest share of data early on.
- *source_agency*: character variable refining the definition of the source by indicating the agency/institution/project from which the data point was obtained: possible values are “iNaturalist”, “PDA” (Pennsylvania Dept. of Agriculture), “NJDA_Public_reporting” (New Jersey Dept. of Agriculture’s Public Reporting tool), “PDA_Public_reporting” (Pennsylvania Dept. of Agriculture’s Public Reporting tool), “DDA” (Delaware Dept. of Agriculture), “ISDA” (Indiana State Dept. of Agriculture), “MDA” (Maryland Dept. of Agriculture), “NYSDAM” (New York State Dept. of Agriculture and Markets), “VDA” (Virginia Department of Agriculture and Consumer Services), “VA_Tech_Coop_Ext” (Virginia Polythecnic and State University/Cooperative Extension), “USDA”.
- *collection_method*: character string defining the method used to collect data: “individual_reporting” for data collected through iNaturalist and public reporting tools, and “field_survey/management” for data collected by agencies in the field. The accuracy of self-reporting data might be lower than that collected by field surveyors.
- *year*: integer value defining the calendar year when the information was collected
- *bio_year*: integer defining the biological year when the information was collected. The biological year follows the species’ development schedule and starts around the time of the emergence of first–instar nymphs (May 1^st^–April 30^th^).
- *latitude*: expressed in decimal degrees (WSG84 coordinate system)
- *longitude*: expressed in decimal degrees (WSG84 coordinate system)
- *state*: character defining the state where the data was collected (two-letter abbreviation, https://www.faa.gov/air_traffic/publications/atpubs/cnt_html/appendix_a.html)
- *lyde_present*: logical value defining whether records are present for spotted lanternfly at the site at the time of survey. These might include regulatory incidents where a single live individual or a small number of dead individuals were observed at the site, but no signs of established population could be detected.
- *lyde_established*: logical value defining whether signs of an established population are present at the site at the time of survey. These include a minimum of 2 alive individuals or the presence of an egg mass as per the working definition of establishment provided by the USDA.
- *lyde_density*: ordinal variable defining the population density of spotted lanternfly at the site, estimated directly as an ordinal category by the data collector or derived from count data. The categories include: “Unpopulated”, indicating the absence of an established population at the site (but not excluding the presence of spotted lanternfly in the form of regulatory incidents); “Low”, indicating an established population is present but at low densities, reflecting at most about 30 individuals or a single egg mass; “Medium”, indicating the population is established and at higher densities, but still at low enough population size to allow for a counting of the individuals during a survey visit (a few hundred at most); “High”, indicating the population is established and thriving, and the area is generally infested, to a degree where a count of individuals would be unfeasible within a survey visit.
- *pointID*: character string uniquely identifying each data point.

### Package installation and data access

The `lydemapr` package can be installed in two different ways. The public repository allows the user to install the package directly from github, by executing the following command in a local R or RStudio instance: `devtools∷install_github(“ieco-lab/lydemapr”)`. This requires the package `devtools` (Wickham et al., 2022) and its dependencies to be installed locally. Alternatively, the package can be obtained by cloning the repository from the github page https://github.com/ieco-lab/lydemapr. The package can then be installed locally by opening the file `lydemapr.Rproj` in RStudio, and clicking “Install package” in the Build tab. Once the package is installed, the user has access to the complete dataset, which can be loaded by typing `lydemapr∷lyde` in the R console. In addition, the package contains a summarized version of the same dataset at a lower spatial resolution (10 km^2^), which can be accessed by typing `lydemapr∷lyde_10k` instead.

If the user is only interested in accessing the data without using the R package, or is unfamiliar with R, all datasets contained in `lydemapr` are available for download through the same github repository page (https://github.com/ieco-lab/lydemapr/blob/main/download_data/lyde_data.zip), where the user can download a compressed file containing the data (in .csv format) and Metadata associated with it. All information concerning package installation and data access is available at the frontpage of the github repository.

### Package functions

For a summary overview of the data, the function `lyde_summary()` provides a breakdown of the dataset, showing the number of data points collected each year in each state where data has been collected. The package contains two customizable functions that can be used to visualize the data spatially. The function `map_spread()` provides an up-to-date map displaying the progression of the established invasion range through time, in addition to the locations of surveys which did not detect established populations (Figure 2). Function arguments allow the user to select the spatial resolution at which the data should be mapped, and the spatial extent of the figure produced. A second function included in the package, `map_yearly()` maps the recorded population density at a 10km^2^ resolution broken down by year (Figure 3). Through this visual depiction, it’s possible to observe where survey efforts have been focusing on each year, as the invasion front progressed.

**Figure 2.**
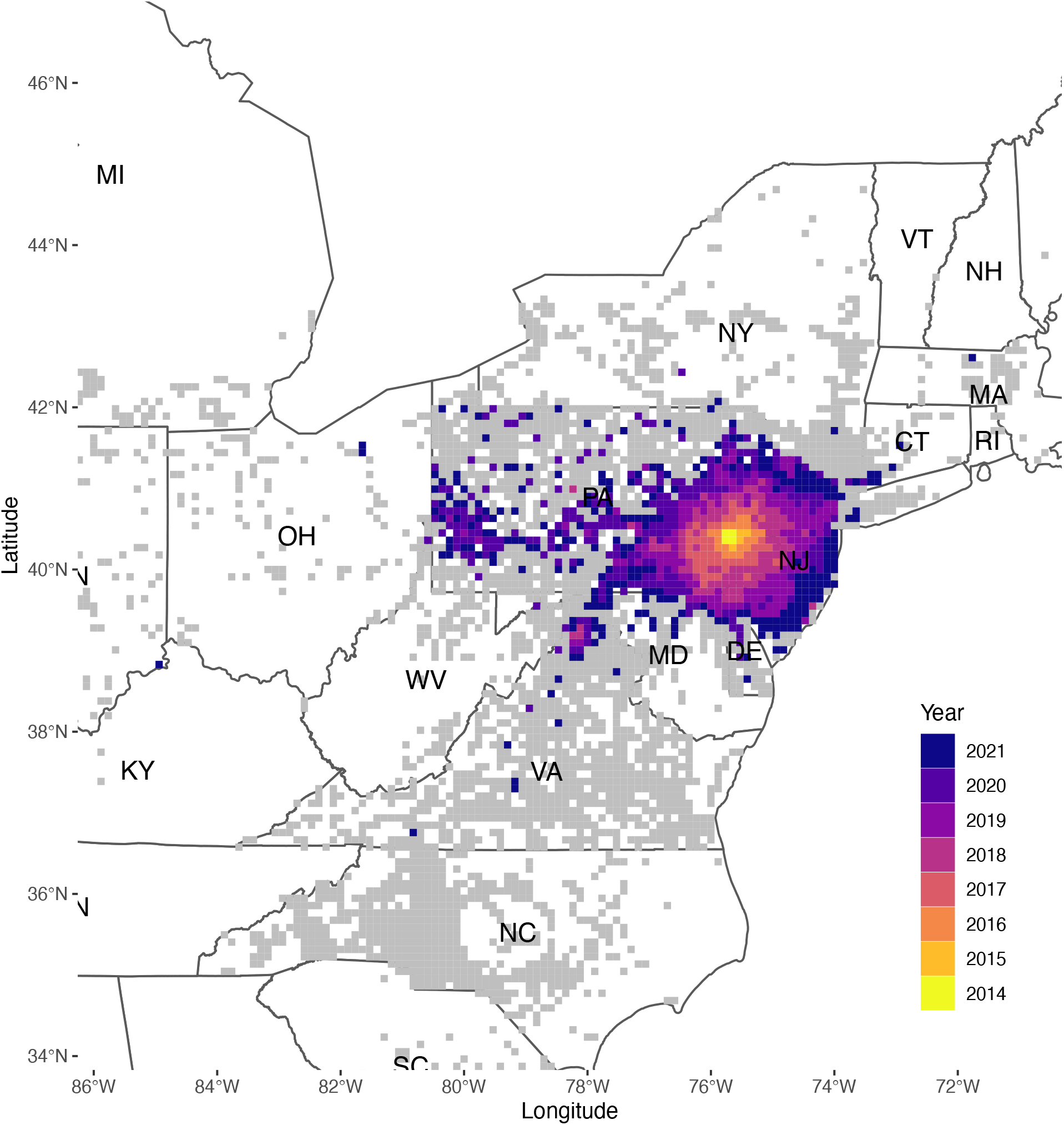
Map produced through the package function `map_spread()`. The map shows the year of first discovery of established populations of the spotted lanternfly (colored tiles) in 10km^2^ grid cells across the Eastern United States, as well as the location of negative survey records for the establishment of the species (grey tiles).

**Figure 3.**
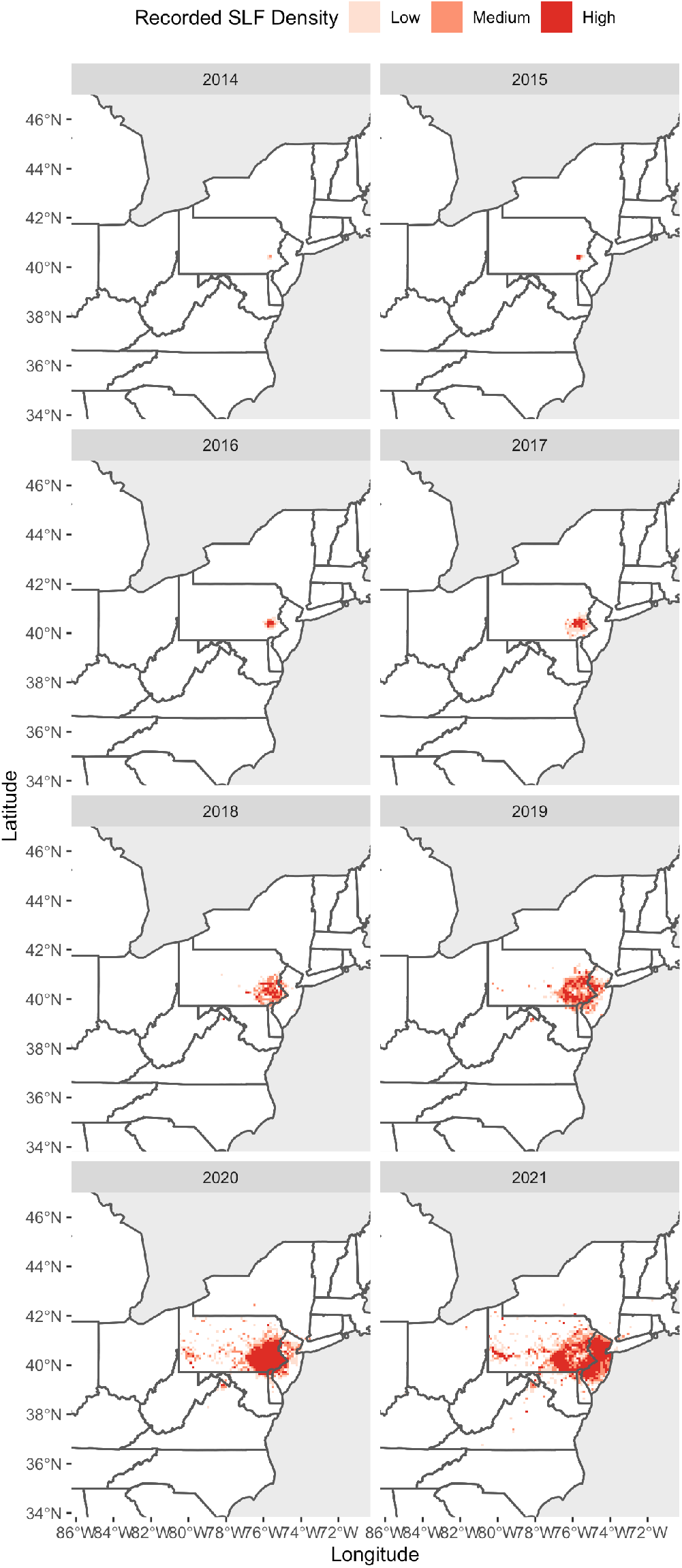
Map produced through the package function `map_yearly()`, showing the population density of spotted lanternfly assessed yearly in 10km^2^ grid cells across the Eastern United States (red tiles).

## Conclusion

Several models have been developed to forecast the future spread and establishment potential of spotted lanternfly in the United States (Huron et al., 2022; Jones et al., 2022; Lewkiewicz et al., 2022; Wakie et al., 2020) and elsewhere (Jung et al., 2017; Maino et al., 2022). To ensure future models can be compared and combined through ensemble procedures, models should be based on the same historic and present spread data of spotted lanternfly. In addition, a comprehensive dataset can provide additional information on population trends through time in specific areas, as well as offer insight on the efficacy of control actions over time.

## Author’s contribution

SDB and MRH conceived the paper, gathered the data, and wrote the code for the package. LB, PK, JL, GRP provided survey data and helped harmonize it across sources. All authors contributed with the writing of the manuscript.

## Acknowledgements

We would like to thank Eric Day for providing data on a citizen science project run by the Virginia Polytechnic Institute and State University and the Virginia Cooperative Extension. We thank Jocelyn Behm, Stefani Cannon, Anna Carlson, Jason Gleditsch, Stephanie Lewkiewicz, Sam Owens, Payton Phillips, and Timothy Swartz for their insightful comments on early drafts. This work was funded by the United States Department of Agriculture Animal and Plant Health Inspection Service Plant Protection and Quarantine under agreements AP19PPQS&T00C251, AP20PPQS&T00C136, AP20PPQS&T00C118, AP22PPQS&T00C146 and AP22PPQS&T00C097; the United States Department of Agriculture National Institute of Food and Agriculture Specialty Crop Research Initiative Coordinated Agricultural Project Award 2019-51181-30014; the Pennsylvania Department of Agriculture under agreements 44176768, 44187342, C9400000036, and C94000833; and the California Department of Food and Agriculture under agreement A20-0850-000-SA.

## Data availability

The package, containing the open access data, is stored as a public repository at https://github.com/ieco-lab/lydemapr.

